# A deep learning and novelty detection framework for rapid phenotyping in high-content screening

**DOI:** 10.1101/134627

**Authors:** Christoph Sommer, Rudolf Hoefler, Matthias Samwer, Daniel W. Gerlich

## Abstract

Supervised machine learning is a powerful and widely used method to analyze high-content screening data. Despite its accuracy, efficiency, and versatility, supervised machine learning has drawbacks, most notably its dependence on *a priori* knowledge of expected phenotypes and time-consuming classifier training. We provide a solution to these limitations with *CellCognition Explorer*, a generic novelty detection and deep learning framework. Application to several large-scale screening data sets on nuclear and mitotic cell morphologies demonstrates that *CellCognition Explorer* enables discovery of rare phenotypes without user training, which has broad implications for improved assay development in high-content screening.

## Introduction

Advances in microscope automation have facilitated the systematic study of cellular phenotypes resulting from genetic or chemical perturbations. Prevailing image analysis pipelines rely on supervised machine learning to classify cellular phenotypes based on user-defined collections of statistical image features (Bakal et al., 2007; Boland and Murphy, 2001; Boutros et al., 2015; Carpenter et al., 2006; Held et al., 2010; Jones et al., 2009; Mattiazzi Usaj et al., 2016; Misselwitz et al.; Neumann et al., 2006; Ramo et al., 2009; Sommer and Gerlich, 2013). This approach has revealed new gene functions in various RNA interference (RNAi)-based screens (e.g., (Boutros et al., 2015; Cuylen et al., 2016; Goshima et al., 2007; Gudjonsson et al., 2012; Liberali et al., 2014; Neumann et al., 2010; Schmitz et al., 2010; Sommer and Gerlich, 2013)). While supervised machine learning is in principle broadly applicable, it requires extensive user interaction for the development of new biological assays.

A major limitation of supervised machine learning is the requirement for classifier training for each new assay or variation in the experimental conditions. The classifier training requires representative images for all expected phenotype classes. As all possible phenotype morphologies are not always completely known *a priori*, screening assay development often involves extensive and time-consuming pilot screens that are inspected visually for manual annotation of phenotype classes (Conrad and Gerlich, 2010). This process can be facilitated by interactive learning (Jones et al., 2009), yet rare phenotype classes might not be represented in pilot screens. Moreover, current high-content screening analysis software relies on user-curated collections of feature extraction algorithms, which often require specific software adaptations when establishing new cell biological assays. Developing classifiers for new biological assays has hence remained a major bottleneck in high-content screening.

These limitations might be overcome by unsupervised learning methods, which estimate phenotypic content based on intrinsic data structure. However, high cell-to-cell variability and experimental noise often preclude sensitive and reliable detection of low-penetrance phenotypes. The accuracy of unsupervised phenotype detection can be improved by object features beyond the single-cell level, as for example the temporal context (Failmezger et al., 2013; Zhong et al., 2012) or the cell population context (Liberali et al., 2014; Rajaram et al., 2012), but such information is not always applicable for a given biological assay. The detection of unknown phenotypes might be facilitated by novelty detection methods (Manning and Shamir, 2014; Yin et al., 2013; Yin et al., 2008), yet the performance of such methods in genome-wide screening has not yet been tested.

The dependency of machine learning approaches on user-curated image features might be overcome by feature self-learning from the data. This can be achieved by deep learning methodology (LeCun et al., 2015), which has demonstrated impressive performance in various domains, as for example face recognition (Taigman et al., 2014) and speech recognition (Sainath et al., 2013). Recent analyses of protein localization in budding yeast by deep learning indicates a high potential in bioimaging (Kraus et al., 2017; Parnamaa and Parts, 2017), but applicability to genome-scale human cell screening data has not yet been explored and an integration of novelty detection with deep learning is not available in community-standard software for high-content screening.

We here present the novelty detection and deep learning software *CellCognition Explorer*. The software yields phenotype scores based on deviation of cell morphologies from negative control images. We demonstrate that *CellCognition Explorer* enables sensitive and accurate cellular phenotype detection in genome-scale screening data without the need for extensive user interaction for data annotation. In addition, deep learning of image-derived features overcomes the dependency on user-curated feature analysis collections and accurate cell segmentation outlines. Hence, *CellCognition Explorer* greatly facilitates rapid screening assay development even when cellular phenotypes are not known *a priori*.

## Results

### CellCognition Explorer software

We have developed *CellCognition Explorer*, a machine learning framework for the detection of abnormal cell morphologies without prior user training. Two core technologies enable the unsupervised classification task: Deep learning methods infer numerical descriptors of individual cell objects directly from the raw image pixel data, thus circumventing feature engineering and software adaptations for new biological assays. Novelty detection methods then learn a statistical model of the natural phenotype variation within the negative control cell population. This yields an accurate classification boundary for subsequent detection of any morphological deviations in large-scale screening data – even for phenotypes that are not known *a priori*.

*CellCognition Explorer* provides a pipeline for integrated data analysis from raw images to phenotype scores (Fig. 1). The software package consists of two programs: the main *CellCognition Explorer* program provides interactive data visualization tools and the possibility to perform versatile analysis workflows using novelty detection methodology, as well as conventional supervised learning methods. It is controlled by a simple graphical user interface (Supplementary Fig. 1) that works on all major computer operating systems. *CellCognition Deep Learning Module* is a separate program for graphics processing unit (GPU) - accelerated high-performance computing of deep learning features (Supplementary Fig. 2). The implementation in two separate programs provides optimal flexibility for installation of the interactive data exploration and workflow design tool, while enabling the efficient computing of deep learning features by dedicated GPU hardware. Both programs are controlled by graphical user interfaces and distributed as open source software embedded within the *CellCognition* platform (Held et al., 2010) (http://software.cellcognition-project.org/explorer/). Inter-operability with widely used bioimage software packages, including ImageJ/Fiji (Abramoff et al., 2004; Schindelin et al., 2012), CellProfiler (Carpenter et al., 2006), and Bioconductor (Gentleman et al., 2004), is enabled via the standardized file format *cellH5* (Sommer et al., 2013).

**Figure 1.**
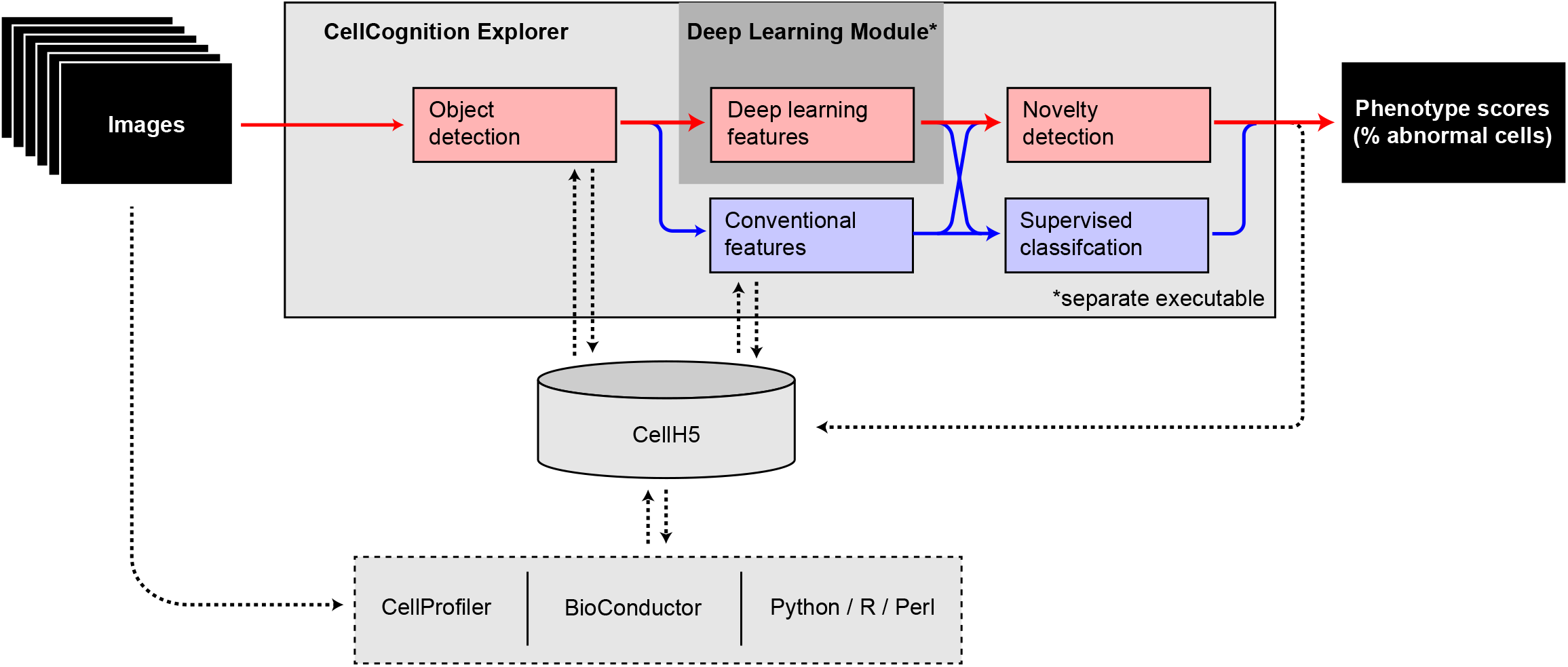
Data analysis workflows with CellCognition Explorer. Red boxes and arrows indicate the standard workflow with object detection, deep learning, and novelty detection as described in this paper. Blue boxes indicate previously described methods (Held et al., 2010) that can be combined with the new functionality (blue arrows) or with other high content screening software via CellH5 (Sommer et al., 2013) (dashed lines). Deep learning features are computed through a separate program (owing to specific high-performance computing hardware requirements). Workflow integration is achieved through CellH5 (Sommer et al., 2013) data exchange.

**Figure 2.**
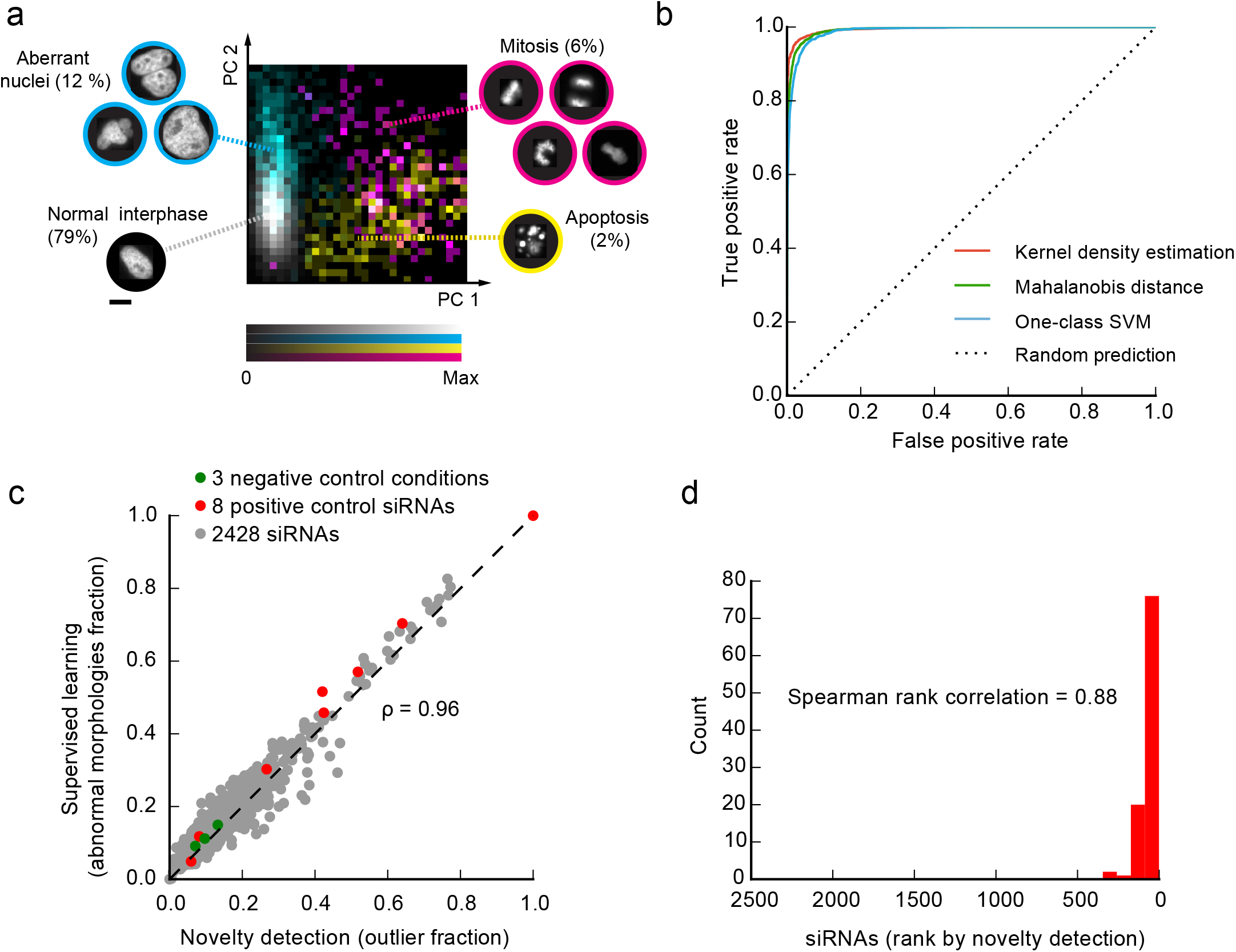
Novelty detection with CellCognition Explorer. (a) Phenotype distribution in principal component sub-space of 10,000 manually annotated HeLa cells stably expressing H2B-mCherry. The cell images were randomly selected from 2,428 RNAi experiments to establish a representative collection of all cellular phenotypes. The 9 phenotype classes illustrate the most common nuclear and mitotic chromatin morphologies. Colors indicate phenotype groups as illustrated by examples. (b) Performance of different novelty detection methods. True positive and false positive rates (receiver operating characteristic curve) were calculated for the indicated methods based on the data shown in (a). (c) Comparison of phenotype scoring by conventional supervised learning (Held et al., 2010) versus novelty detection. Each dot represents one of 2,428 different siRNAs; negative controls: non-targeting siRNAs; positive controls: siRNAs causing strong mitotic phenotypes. Phenotype scores are defined as the fraction of HeLa-H2B-mCherry cells deviating from normal interphase morphology. For detailed analysis results, see Supplementary Table 2. (d) Phenotype scoring of 2,428 siRNAs as in (a) by novelty detection using CellCognition Explorer. Red bars indicate the distribution of the top-100-ranked siRNA hits identified by conventional supervised learning, see (c).

### Cell phenotyping by novelty detection

The novelty detection algorithms implemented in *CellCognition Explorer* are designed to autonomously learn intrinsic cell-to-cell variability in an untreated negative control cell population, which sensitizes the classifier towards perturbation-induced phenotypes. Abnormal cell phenotypes are then scored either based on the weighted cell object distance in feature space relative to the mean and covariance of a control cell population (Mahalanobis distance, MD (Pimentel et al., 2014)), based on their likelihood after estimating the multi-variate probability density of control conditions (kernel density estimation, KDE (Pimentel et al., 2014)), or by fitting a non-linear hyper-plane to control cell objects (one-class support vector machine (Scholkopf et al., 2001)) (see Materials and Methods). These methods can score and classify cell objects based on conventional pre-computed numerical features (e.g., (Boland and Murphy, 2001; Carpenter et al., 2006; Held et al., 2010; Murphy et al., 2003)) or based on learned representations from the original image pixel data when combined with deep learning (Durr and Sick, 2016; Kraus et al., 2016; Vincent et al., 2010).

To test the performance of the novelty detection methods, we generated a data set representing the full spectrum of morphology phenotypes for a chromatin marker. HeLa cells stably expressing fluorescent histone 2B fused to mCherry (H2B-mCherry) were subjected to RNAi depletion of 1,214 genes previously identified as important for mitosis from a genome-wide screen (Neumann et al., 2010), using individual transfection of two different small interfering RNAs (siRNAs) per target gene, followed by live cell imaging on an automated epifluorescence microscope. To establish a reference annotation, we detected cell objects based on local adaptive thresholding and watershed segmentation and calculated 239 conventional numerical features describing texture and shape (Held et al., 2010). We then randomly subsampled 10,000 cell objects and manually annotated their “ground truth” phenotypes by classification into 9 different morphology classes: normal interphase nuclei and various different outlier morphologies, representing mitotic stages, dead cells, and abnormal nuclear shapes (Fig. 2a, Supplementary Table 1, full data set available at http://software.cellcognition-project.org/explorer/). In the principal component subfeature space (Pimentel et al., 2014), interphase nuclei appeared as a single compact cluster, whereas the different outlier phenotype groups scattered broadly in different regions.

All three novelty detection methods implemented in *CellCognition Explorer* accurately classified normal interphase nuclei as inlier objects and other morphologies as outliers, consistent with phenotype scoring with supervised analysis (Fig. 2b). To extend the performance tests to the full data set, we next quantified the abundance of abnormal cell phenotypes in each of the 2,428 RNAi conditions. The fraction of cells with outlier morphologies calculated with novelty detection methods consistently matched the reference state-of-the-art analysis using supervised learning by support vector machines (Held et al., 2010) (SVM, Fig. 2c, d, and Supplementary Fig. 3). Thus, novelty detection with *CellCognition Explorer* accurately identifies abnormal cell morphology phenotypes without the need of extensive data annotation.

**Figure 3.**
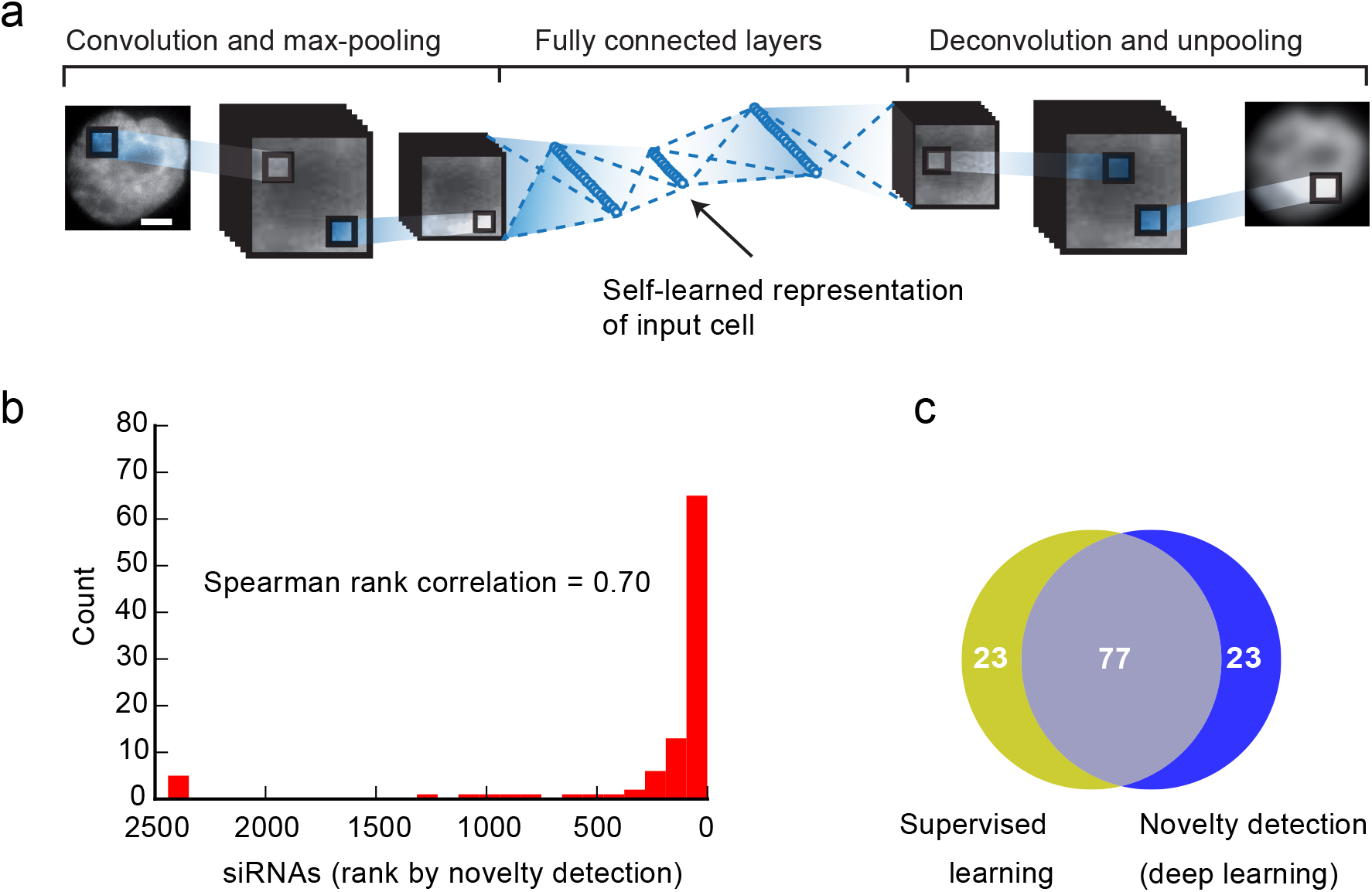
Self-learning of cell object features with CellCognition Deep Learning Module. (a) Schematic illustration of deep learning using an autoencoder with convolutional, pooling, and fully connected layers. (b) Phenotype scoring of 2,428 siRNAs (see Fig. 1a) by novelty detection and deep learning using CellCognition Explorer. Red bars indicate the distribution of the top-100-ranked siRNA hits identified by conventional supervised learning as in (Held et al., 2010). (c) Comparison of the top-100 screening hits determined either by novelty detection and deep learning of object features (blue) or supervised learning and conventional features (yellow) for 2,428 siRNAs as in (a, b). For comparison of novelty detection with conventional and deep learning features, see Supplementary Figure 4a. Scale bars, 10 µm.

### Deep learning of cell features

*CellCognition Deep Learning* Module automatically extracts numerical feature sets that adjust to specific cell morphology markers used in an assay. This is achieved by a convolutional autoencoder, a multilayered artificial neural network (Hinton and Salakhutdinov, 2006) that learns a representation (encoding) for a collection of images. This method requires only center coordinates of cell objects as an input and is thus independent of accurate object segmentation contours that are normally necessary to calculate shape features by conventional user-curated feature sets. The features derived by deep learning serve as an input for the novelty detection method (see above) and can also be used for conventional supervised machine learning. The implementation of different analysis pipelines combining supervised and unsupervised methods is facilitated by the interactive visualization platform for cell objects (Fig. 1).

We evaluated the accuracy of phenotype scoring based on deep learning-derived features using the reference data as described above. We trained deep autoencoder neural networks to reduce the highdimensional image-pixel data to a low-dimensional compressed code (Fig. 3a, for details see Supplementary Tables 5-7). The parameters of such networks are iteratively adjusted by minimizing the discrepancy between the original data and its reconstruction based on the compressed code. The learned features then serve as an input for novelty detection as described above. The top-scoring RNAi phenotypes obtained by this method matched well to the reference scoring by supervised learning (Fig. 3b, c). The total accuracy achieved by deep learning features was only slightly lower than the classical feature collection (compare Supplementary Fig. 4b, c), which has been highly optimized towards the specific chromatin morphology assay. Thus, unsupervised deep learning can derive informative features for fully automated phenotype scoring by novelty detection, thereby overcoming the dependency on manually curated feature sets.

**Figure 4.**
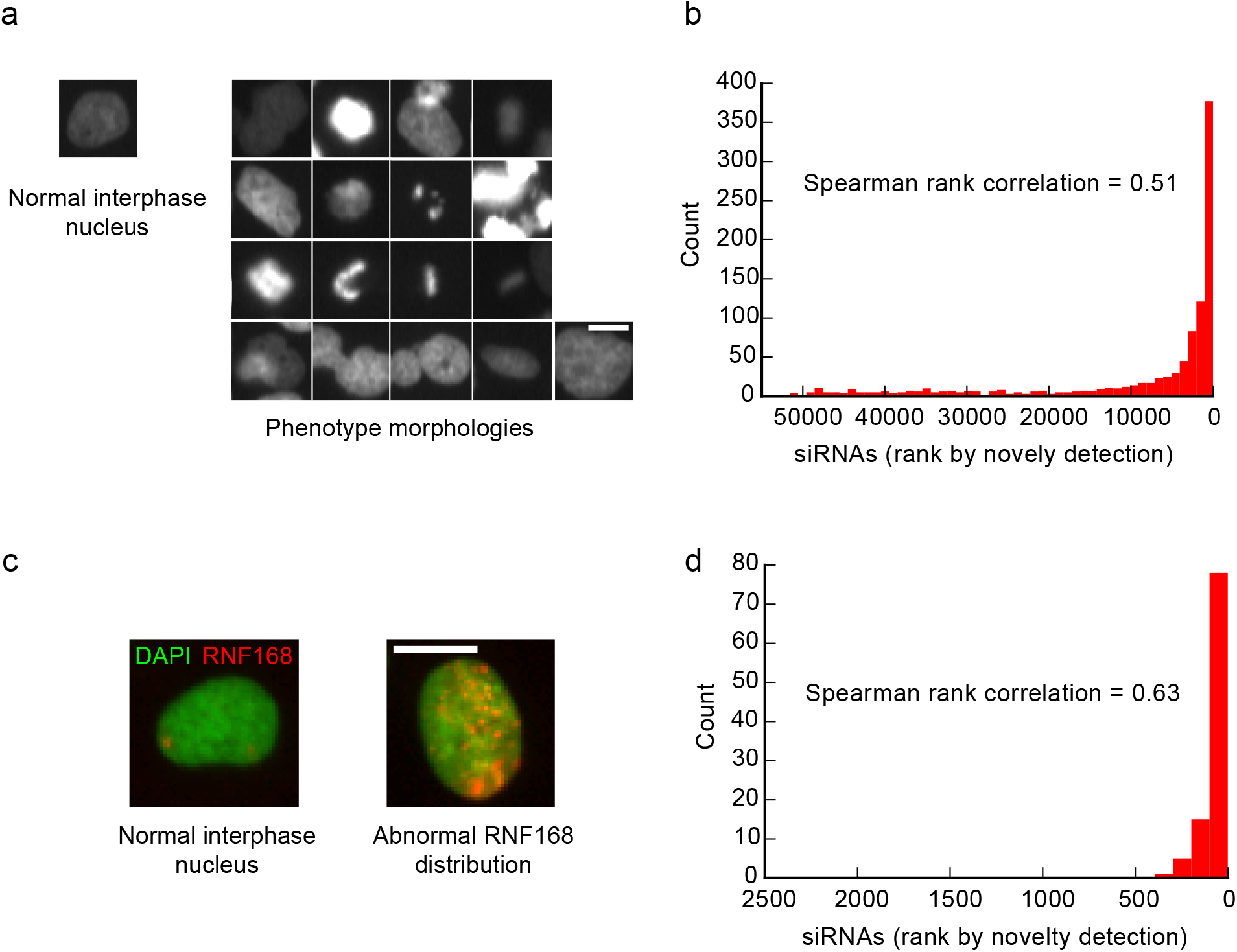
Application of CellCognition Explorer to high-throughput RNAi screening. (a) Genome-wide RNAi screen for mitotic regulators based on live-cell microscopy of HeLa cells expressing H2B-mCherry (Neumann et al., 2010). Images show representative examples for a normal interphase morphology and various phenotype morphology classes as defined in (Neumann et al., 2010). (b) Phenotype scoring of 51,748 siRNAs by novelty detection and deep learning with CellCognition Explorer, using the primary image data of (Neumann et al., 2010). Red bars indicate the distribution of the top-1000-ranked siRNA hits identified by conventional supervised learning as in (Neumann et al., 2010). For details on analysis, see Supplementary Table 3. (c) Large-scale RNAi screen for DNA damage repair regulators (Gudjonsson et al., 2012). GFP-tagged RNF168 (red) visualizes spontaneous DNA damage in U2OS cells by accumulation in bright nuclear foci, which increase in number and intensity upon perturbed DNA damage repair (Abnormal RNF168 distribution in siTRIP12 RNAi cell). DNA (green) is counterstained by DAPI. (d) Phenotype scoring of 2,423 siRNAs by novelty detection and deep learning with CellCognition Explorer, using the primary image data of (Gudjonsson et al., 2012). Red bars indicate the distribution of the top-100-ranked siRNA hits as identified in (Gudjonsson et al., 2012). For details on analysis, see Supplementary Table 4. Scale bars, 10 µm.

### Application to genome-wide screening

To test the applicability of *CellCognition Explorer* to high-throughput screening, we aimed to re-compute phenotype scores for published primary image data of a genome-wide RNAi screen for mitotic regulators (Neumann et al., 2010). The original phenotype scoring of these screening data was based on classification using supervised learning (Neumann et al., 2010), whereby the development of a reliable classifier required a multi-month phase of iterative pilot screening and visual inspection for classifier training (Neumann et al., 2006; Neumann et al., 2010). The analysis of millions of images during pilot screening had ultimately led to the discovery of many previously unknown phenotypes (Fig. 4a). Identifying all abnormal cell morphologies prior to classifier training would abrogate the need to visually inspect the full dataset, thereby solving a major bottleneck in data analysis.

We used the *CellCognition Deep Learning Module* to extract self-learned features for phenotype scoring based on novelty detection. The top-ranked siRNAs according to phenotype penetrance were very similar to the originally published hit list (Fig. 4b), even though information on abnormal phenotype morphologies from pilot screening was not taken into account. The generation of hit-lists by our new methods are hence highly enriched for abnormal cell morphologies, which greatly facilitates further phenotype analysis by visual inspection, supervised classification, or unsupervised clustering.

To probe the versatility of our methods, we next re-analyzed primary image data from another large-scale RNAi screen for DNA damage repair regulators (Gudjonsson et al., 2012). The primary assay probed the regulation of DNA damage repair protein RNF168, which accumulates in bright nuclear foci (Fig. 4c). The distribution of top-ranked siRNA hits derived from deep learned-features and novelty detection was very similar to the original conventional supervised learning analysis (Fig. 4d). Hence, *CellCognition Explorer* yields reliable phenotype scores in large-scale screening data without prior user definition of aberrant phenotype morphologies.

To ensure broad applicability in biological laboratories, we designed our software to achieve high-throughput data processing independent of expensive computing infrastructure. We therefore implemented the computationally intense deep learning module as a separate software that runs on standard consumer-grade graphics hardware on a desktop computer (see methods).

With this, we completely processed both large-scale screening datasets. The genome-wide RNAi screen (Fig. 4a, b) was processed in less than 34 h in total. The autoencoder training on 200,000 cell objects from negative control conditions took 2 h. After training, 19 million cells were classified with the novelty detection procedure (including feature generation) in 32 h, hence with a throughput of 593,750 cells / h. Thus, deep learning and novelty detection with *CellCognition Explorer* provides a versatile and powerful solution for rapid phenotype scoring in large-scale high-content screening.

## Discussion and outlook

*CellCognition Explorer* bypasses the need for extensive pilot screening to capture all possible phenotype morphologies and thereby solves a major bottleneck of conventional supervised approaches. *CellCognition Explorer* yields a list of top-ranked screening targets directly from large-scale screening data, which can be used for mechanistic follow-up studies or further sub-classified using conventional supervised methods.

By avoiding human bias arising from supervised classifier training (Zhong et al., 2012), novelty detection with *CellCognition Explorer* helps to improve the consistency between different screens. Furthermore, the independence from manually curated feature sets facilitates the development of new cell biological assays. The inference of image features by deep learning does not take the image segmentation contours into account and therefore has the potential to facilitate the analysis of cellular markers that are difficult to segment.

We here demonstrate how *CellCognition Explorer* can be used to detect novel phenotypes in nuclear morphology screening data. *CellCognition Explorer* supports processing of multi-channel images and we are currently extending the software for segmentation of other cell compartments and cell tracking for kinetic readouts (Hoefler, unpublished results).

Overall, the simple graphical user interface of *CellCognition Explorer* and the independence of centralized large-scale computing infrastructure is well suited to apply deep learning and novelty detection to diverse cell biological questions. Hence, *CellCognition Explorer* provides new opportunities for high-content screening as well as exploratory research in diverse biological fields.

## Materials and Methods

### Cell culture

A HeLa cell line stably expressed histone H2B fused to mCherry and lamin B1 fused to EGFP (Daigle et al., 2001) was generated from a HeLa Kyoto cell line as previously described (Schmitz and Gerlich, 2009). HeLa cells were cultured in Dulbecco’s modified Eagle medium (DMEM; Gibco) supplemented with 10% (v/v) fetal bovine serum (FBS; Gibco), 1% (v/v) penicillin-streptomycin (Sigma-Aldrich), 500µg ml^−1^ G418 (Gibco) and 0.5 µg ml^−1^ puromycin (Calbiochem). Homogeneous expression levels of the two transgenes were ensured by fluorescence-activated cell sorting. The cell line was tested negatively for mycoplasm contamination in our quarterly testing routine. The parental HeLa cell line (‘Kyoto strain’) was obtained from S. Narumiya (Kyoto University, Japan) and validated by a Multiplex human Cell line Authentication test (MCA).

### siRNA library transfection

1,114 genes of the MitoCheck genome-wide RNAi screen validation data set (Neumann et al., 2010) were targeted by either of two siRNAs. To test target specificity, all siRNAs were mapped against the 2013 human genome (ENSEMBL V70) resulting in a unique match. siRNAs were delivered in 384-well imaging plates (Falcon) using solid-phase reverse transfection (Erfle et al., 2008). Cells were seeded on the imaging plates using a Multidrop Reagent Dispenser (Thermo Scientific).

### Automated microscopy

40 h after cell seeding, plates were imaged on a Molecular Devices ImageXpressMicro XL screening microscope using a × 20, 0.75 NA, S Fluor dry objective (Nikon). 4 positions with 2 Z-sections each (4 µm offset) were acquired in each well. Images were flat-field-corrected with the Metamorph software (Molecular Devices) using images acquired in empty wells to compensate for inhomogeneous illumination.

### Image pre-processing for reference annotation with conventional supervised learning

To detect individual cells, we used local adaptive thresholding followed by a watershed split-and-merge segmentation as provided in the CellCognition framework (Held et al., 2010). For each cell, the segmentation outline, its center-of-mass, and its bounding-box was computed and saved to the cellH5 format (Sommer et al., 2013). Then, 239 cellular morphology features describing the texture, shape and intensity distribution of each single cell object were computed as in (Held et al., 2010). As reference annotation in the chromatin morphology screen, we trained a multi-class support vector machine on 9 phenotype classes with 592 training examples, achieving a cross-validation accuracy of 85.1%.

### Outlier detection

An outlier is an observation that deviates so much from other observations to arouse suspicions that it was generated by a different process (Hawkins, 1980). Novelty detection methods aim to identify outliers by inference based on intrinsic properties of the data without any further annotations. We implemented three state-of-the-art novelty detection methods: Mahalanobis distance (Mahalanobis, 1936), kernel density estimation (Pimentel et al., 2014), and one-class support vector machine (Scholkopf et al., 2001). All methods find novelty based on pre-extracted morphology features or on data-adaptive, learned feature representations by deep learning autoencoders.

### Mahalanobis distance

The Mahalanobis distance (MD) is based on the parametric assumption that the extracted features follow a multi-variate Gaussian distribution (Mahalanobis, 1936). The Mahalanobis distance (eq. 1) incorporates the mean µ and the covariance matrix **S** (multi-dimensional spread) of the data.

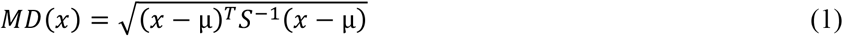

For the estimation of the mean and covariance, data samples were drawn from negative control conditions. Negative control conditions will also contain aberrant phenotypes (e.g. polypoid cells). To reduce the effect of strong outliers in the estimation process, some instances were excluded based on their univariate feature distribution (i.e. cells with the most extreme single feature values) prior to the estimation of the multi-variate mean and covariance.

The mean feature vector represents the ideal normal cell, whereas the covariance estimates its multivariate spread. Once the mean and the covariance of the multi-variate Gaussian distributions are estimated, an empirical cut-off distance is estimated (relative to negative control data) in a way that 15% of the data points have a greater Mahalanobis distance than the chosen cutoff and therefore are considered abnormal. Cells from other experimental conditions are then assigned the normal category (inlier) when their Mahalanobis distance is smaller than the calculated cutoff, or abnormal (outlier) if the distance is greater. For all experiments shown in the manuscript, >50,000 cells were sampled from negative control conditions for estimation the parameters outlier detection. For the chromatin morphology screen (Fig. 2c, d), 10% cut-off was used. Negative control conditions in the screen for damage repair regulators (Fig. 4c, d) contained only very few outliers; a 5% cut-off value was hence chosen. The varying thresholds were empirically determined based on the expected number of outliers in the data-sets.

### Kernel density estimation

Kernel density estimation (KDE) is non-parametric estimator of a probability distribution (Pimentel et al., 2014). Conceptually, KDE is related to histograms. Instead of quantizing a discrete probability density by counting how many data points fall into each bin, KDE does not require binning and outputs a continuous function. KDE places a zero-centered kernel function *K* of a bandwidth *h* on each data point followed by summing all these kernels (eq. 2).

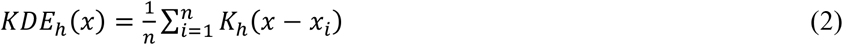

A multi-variate Gaussian density distribution was used as kernel *K*. The bandwidth *h* is given by the variance of the Gaussian distribution. In places where many data points are located, the estimated density will be high, while at places where no or few data points lie, the density will approach zero. Similar to the cut-off parameter of Mahalanobis distance, a separation into abnormal (outlier) and normal (inlier) locations was achieved by thresholding such that the resulting estimated density had 15% of the data points classified novel and 85% classified normal. The choice of an appropriate bandwidth is key to kernel density estimation, since it determines he smoothness of the resulting density (e.g. too small bandwidths lead to spiky non-overlapping densities). Therefore, a cross-validation strategy was used to estimate appropriate bandwidth from hold-out data.

### One-class support vector machine

Another non-parametric and non-linear way to estimate densities in high-dimensional spaces is the one-class support vector machine (OC-SVM) (Scholkopf et al., 2001). Similar to the standard two-class support vector machine (Vapnik and Lerner, 1963) the OC-SVM yields a separating hyper-plane with a maximal margin to distinguish the majority of the data from the origin. When using a Gaussian kernel in the SVM optimization framework, the hyper-plane implicitly corresponds to non-linear decision boundaries in input space (Boser et al., 1992). With this, the OC-SVM models arbitrary densities in high-dimensions as opposed to more simplified ellipsoidal distributions, when using Mahalanobis distance. Similar to the kernel density estimation, a parameter *g* (inverse standard deviation of the Gaussian kernel) accounts for the smoothness of the resulting enclosing hyper-plane. In the experiments shown in the manuscript, *g* was optimized empirically by a grid-search on a logarithmic scale. The parameter *n* in OC-SVM-based novelty detection is the maximal fraction of data points in the training-set, which can be assigned being an outlier. In accordance with the methods described before, we set n=0.15.

### Self-learned feature representation with autoencoders

The novelty detection algorithms described above can be trained on a manually curated feature set (e.g., as in (Held et al., 2010)). Such features are defined *a priori* and do not depend on the actual data at hand. Hence, information for classifying novel phenotypes from normal morphologies might not be sufficiently expressed by general feature sets.

To overcome this limitation, we implemented a class of artificial neural networks (ANNs) termed autoencoders (AE) to learn a data-dependent features representation directly from pixel data from individual cropped-out cells in an unsupervised fashion. The bounding-box size for cropping can be freely chosen by the user. An autoencoder is an artificial neural network, which adjusts its internal weights to reconstruct its input as close as possible (subject to a given objective function) by first learning how to effectively encode its inputs in the encoding layer and then decoding this back to the original domain with an inverse decoding layer (Hinton and Salakhutdinov, 2006). Once an autoencoder has learned how to encode cell images into a code, this process is iterated to stack autoencoders in a hierarchical fashion to generate deeper networks called stacked autoencoders (Hinton and Salakhutdinov, 2006). To ensure that an autoencoder is not learning the simple identity function, the internal parameters have less degrees of freedom than its input / output (contractive autoencoder).

Recent progress in designing deep learning networks for various tasks in computer vision has led to the incorporation of *convolutional* and *pooling layers* (e.g. (Krizhevsky et al., 2012)). Convolutional layers refer to learnable image filters usually of size 3x3 or 5x5 applied to the input images. As other network weights in an ANN these filters are simultaneously adapted during optimization to yield feature-maps, which improve the overall network performance. To reduce the overall spatial dimensionality, convolutional layers are combined with max-pooling layers, which effectively down-samples the input image by selecting only the maximum pixel value for a defined neighborhood region (Krizhevsky et al., 2012)-in our experiments 2x2. We combined the stacked-contractive-autoencoder model with convolutional and pooling layers to achieve more expressive codes in the autoencoder paradigm. For faster autoencoder training *dropout layers* have been proposed (Vincent et al., 2010), which randomly set a fixed percentage of inputs in each layer to zero. This enforces the network to learn to reconstruct from corrupted inputs, which leads to less over-fitting and faster training convergence.

To analyze the data shown in the manuscript, we used autoencoders with a variable sequence of one or more convolutional layers (Supplementary Tables 5-7), followed by max-pooling layers and a central stacked autoencoder. The convolutional and pooling layers are mirrored in the decoding part of the autoencoder with their corresponding inverse de-convolutional and un-pooling layers. Between each layer in the encoding part of the autoencoder, optional dropout layers are inserted with a dropout probability ranging from 10 to 90%. The convolutional and fully-connected layers require a non-linear activation function. In our implementation, *sigmoid (s)* and *linear-rectifier (r)* activation functions are supported.

### Autoencoder layout

To analyze the screening data - chromatin morphology (Fig. 3b, c), MitoCheck RNAi screen (Fig. 4a, b), and DNA damage repair RNAi screen (Fig. 4c, d) - we designed autoencoders such that an input image of a typical cropped single cell (40x40 pixel) is encoded through a series of convolutional layers, pooling layers and fully-connected layers, which successively reduce the input dimensions from 40x40 = 3600 image pixels to a code length of 144 or 64. Note, that the *decoding* part in each autoencoder is composed of layer-wise inversions of the encoding part. The layout of all used autoencoders with precise information about the convolutional and dense layers can be found in Supplementary Tables 5-7.

### Autoencoder training

Autoencoders were trained with stochastic gradient descent (SGD) based on the sum of squared residuals of input and output as objective function (Vincent et al., 2010). The stochastic gradient descent method resembles a stochastic approximation of the gradient descent optimization method, where the true gradient of all the data is approximated by the gradients of so called *mini-batches*, a randomly selected subset of the training data of a certain size *m*. All weights in the network are updated according to the back-propagated gradients of the objective function. This process is repeated and each round is referred to one training epoch. Empirically, we found that alternating two variants of SGD – Nesterov momentum (Nesterov, 1983) and adaptive gradient descent (AdaGrad (Duchi et al., 2011)) - leads to fast convergence. Nesterov momentum adds a momentum term to the update rule to stabilize the learning process and to avoid local minima (Bengio et al., 2013). AdaGrad modifies SGD by introducing a per-parameter learning rate leading to faster convergence (Duchi et al., 2011).

For the autoencoder used in chromatin morphology screen, we used first Nesterov momentum SGD updated for 128 epochs with a mini-batch size of 128, learning rate of 0.02, and a momentum of 0.8 followed by AdaGrad updates for 128 epochs with mini-batch size of 128 and base learning rate of 0.1.

For the autoencoder depicted in MitoCheck RNAi screen, we used a mini-batch size of 128. Nesterov momentum updates with learning rate of 0.1 and momentum of 0.5 were followed by AdaGrad updates with learning rate 0.1, both for 64 epochs. Then, we reiterated with slightly adjusted values: 0.02 learning rate and momentum 0.9 for the Nesterov updates and a learning rate of 0.05 for AdaGrad.

For the autoencoder shown in DNA damage repair RNAi screen, we used first Nesterov momentum SGD with a learning rate of 0.02, momentum of 0.9, and a mini-batch size of 128 followed by AdaGrad with learning rate of 0.05 and a mini-batch size of 64.

### CellCognition Explorer software package

The CellCognition Explorer software package consists of two programs. 1) The CellCognition Explorer main program, from which the novelty detection methods (e.g. Mahalanobis distance) and supervised learning algorithms (multi-class support vector machine) can be executed and visualized; 2) A program for the generation of self-learned feature representations (codes).

#### *CellCognition Explorer* **main program**

The software imports images of cell populations and converts them into galleries of individual cells. Original multi-channel raw images are preprocessed in advance i.e. thumbnails, features and outlines are calculated and saved to cellH5/hdf5. The software stores experimental conditions along with cell objects to enable efficient data export and visualization of results by statistical plots, such as object class counts per condition. The software provides functions for cell segmentation and conventional feature extraction as in (Held et al., 2010) for multiple image channels.

The graphical user interface is designed to handle a few thousand cells smoothly on a standard desktop PC. Larger data sets can be loaded in batches. The software is available as a binary standalone version for Mac OSX and Windows 7.

#### CellCognition Explorer Deep Learning Module

With this program, the user can train a deep learning autoencoder from negative controls. The main input is a cellH5 file (Sommer et al., 2013) containing the image data and center positions of all detected cell objects, together with a position mapping (text-)file indicating negative control positions used for randomly sampling cells for the training process. After training, the resulting autoencoder model can be used in the *encoding* step. The output file of the encoding step can directly be loaded into the *CellCognition Explorer* main program for novelty detection and / or supervised learning. *CellCognition Explorer Deep Learning Module* provides a graphical user interface to specify all processing parameters (Supplementary Fig. 2). The software is provided as source code (LGPL) and as binary installer using the docker framework. For installation in MacOS X, a docker execution script is provided.

### Implementation details

CellCognition Explorer is implemented in Python 2.7 and the deep learning module makes use of the Theano (ver. 0.7.0) and lasagna (ver. 0.2) library for deep learning and GPU computing interfacing with NVIDIA CUDA (ver. 5.5). For the graphical user interfaces, PyQt5 (ver. 5.3.2) binding to the C++ library Qt (ver. 5.3.1) was used. Novelty detection methods are based on the sklearn library (ver. 0.16.0). Image-based screening data is segmented and processed using CellCognition Analyzer (ver. 1.6.0) and analysis results were saved in the cellH5 format (ver. 1.3.1).

### License

CellCognition Explorer is released under GNU General Public License version 3 (GPLv3).

### Computing hardware

All data shown in this manuscript was processed on a HP-Z820 desktop workstation with Intel^®^ Xeon^®^ CPU E5-2665-0 @ 2.40 GHz (16 CPUs) equipped with 64 GB of RAM and a NVIDIA GeForce GTX TITAN X graphics adapter.

## Acknowledgments

D.W.G. has received funding from the European Community’s Seventh Framework Programme FP7/2007-2013 under grant agreement no. 241548 (MitoSys) and no. 258068 (Systems Microscopy), an ERC Starting Grant under agreement no. 281198 (DIVIMAGE), and from the Austrian Science Fund (FWF) project no. SFB F34-06 (Chromosome Dynamics). The authors thank the IMBA/IMP BioOptics facility for technical support, Claudia Lukas, Thomas Walter, Jean-Karim Hériché, Beate Neumann, and Jan Ellenberg for providing original image data of published RNAi screens, Thomas Walter for assistance with image pre-processing, Sara Cuylen for assistance with the graphical user interface design, and Life Science Editors for editorial support.

## Author contributions

Conceived the project: D.W.G., C.S.; designed experiments: D.W.G., M.S.; performed experiments: M.S.; analyzed data: C.S.; implemented software: R.H. *(CellCognition Explorer* main program); C.S. *(CellCognition Deep Learning Module);* wrote the paper: D.W.G., C.S.

## Competing financial interests

The authors declare no competing financial interests.

